# Analyses of Genome-Wide Recombination in *Sulfolobus islandicus*

**DOI:** 10.64898/2025.12.16.694729

**Authors:** David J. Krause, Changyi Zhang, Rachel J. Whitaker

## Abstract

Recombination is a fundamental process that both maintains genome integrity and dramatically shapes evolutionary dynamics in populations from all domains of life. Here we cross two genetically distinct but highly syntenic sympatric strains of *Sulfolobus (Saccharolobus) islandicus* and identify the pattern of recombination across the archaeal chromosome. We quantify recombination breakpoints occurring at neutral chromosomal markers through high-throughput short read Illumina sequencing of selected recombinant strains. We focus on three datasets that provide different, but corroborating information: i) individual colony sequences, ii) large colony pools of strains sorted by background genotype, and iii) a large colony pool of combined background genotypes. For each dataset we apply a newly designed and optimized computational tool called MARBL (<u>M</u>ultiply <u>A</u>nchored <u>R</u>ecombination <u>B</u>reakpoint <u>L</u>ocator) for quantifying breakpoints and testing their statistical significance relative to sources of error introduced by sequencing technology. By combining the results of each dataset, we determine the short tract recombination preference in this organism and discuss its implications for mechanisms of recombination in archaea. We demonstrate uniform rates of recombination throughout the chromosome, which differ from previous studies of natural variation and discuss this in light of previous data suggesting speciation occurs through selection against inter-species hybrids.

**Importance Statement:** Recombination is a critical process in all domains of life that facilitates the essential repair of chromosomes and exchanges variation between organisms. Here we investigate genome-wide patterns of recombination in the model archaeaon *Saccharolobus islandicus* to better understand mechanistic aspects of their DNA exchange and recombination processes. Our observations of recombination in a controlled laboratory setting identified patterns of recombination that support the ability to incorporate short pieces of DNA in a piecemeal fashion from larger transferred segments. By comparing these observations to inferred recombination from natural populations, we can gain insight into evolutionary parameters including natural selection that would be otherwise difficult to determine from genomic evidence alone.

## Introduction

Recombination rate variation modulates the effect of selection on genomic diversity, speciation, genome architecture and genome stability (1). Apparent recombination rates can be estimated in natural populations from patterns of genetic variation (2-4). Such studies have yielded tremendous insight into the impact and mechanisms of recombination rate variation in sexual eukaryotes, including reduced levels of crossing over near centromeres (5), sequence-motif associated recombination hotspots (6), and many other cases of chromosome-scale variation in recombination rates (7-9). Studying recombination in natural populations of bacteria and archaea has been enabled by access to genomic data through advances in sequencing technology (10-13).

Standard DNA exchange protocols with selectable markers between genetically diverse strains has yielded strategies for studying genome-wide recombination in the laboratory without environmental selection. Most experimental mechanisms for establishing recombination events and their signatures in genomes conflate mechanistic effects of DNA transfer and homologous recombination. In organisms where either transfer mechanism or properties of homologous recombination have been established, the two processes can be deconvoluted. For example in bacteria and archaea, genetically diverse strains can be crossed and recombinant progeny recovered using selectable markers, and although this type of procedure involves a round of selection, the selection occurs on a well-defined drug resistance or prototrophy-conferring marker gene (17-21). Most commonly for eukaryotes, this type of analysis can be performed by analyzing meiotic products, rather than successful fertilized progeny in humans, flies, and yeast, generating inherent meiotic recombination frequency maps (14-16).

Gene transfer mechanisms described for archaea are diverse and mostly considered to be similar to bacterial conjugation, transformation, and transduction. In the Archaea, the exchange of genetic markers can be observed in *S. islandicus* as well as the related species *S. acidocaldarius* and *S. solfataricus* (21,22). One mechanism proposed for the exchange of DNA occurs via a process mediated by physical contact, suggested by increases in cellular aggregation and gene transfer following exposure to UV-irradiation or DNA-damaging agents (23-25). Some proteins involved in DNA transport between cells have been identified, giving some insight into DNA transport mechanisms in these archaea (26). Recent experiments in *S. islandicus* have found that plasmids can mediate the process of genetic exchange between cells promoting high frequency of recombination on one side of an integrative conjugative plasmid. At some low frequency, this study also found large chromosomal regions distal to the plasmid were also transferred.(27).

The size of individual DNA fragments during genetic exchange sets an upper bound on recombinant tract length, while the mechanism of downstream genetic recombination may influence whether the exchanged DNA is incorporated as one large piece or is broken into many. In the case of transformation, the upper limit of DNA size is likely capped at the limits of the DNA uptake machinery. Plasmid-mediated chromosomal transfer is likely limited by the stability of mating bridges. In cell-cell fusions, the entire chromosome would be accessible, similar to eukaryotic meiosis. Gene transfer experiments using genetically engineered versions of naturally diverse strains can provide single base-pair resolution of genome-wide recombination. In transformations of *Haemophilus influenzae* and *Streptococcus pneumoniae*, studies have identified genome-wide incorporation of donor DNA with recombination tract lengths of several Kb (17,28). In conjugation experiments of *Streptococcus agalactiae* and *Mycobacterium smegmatis*, where integrated plasmids mobilize large chromosomal regions, recombination occurred in tract lengths of 30-120Kb (18,29). In cell-cell fusions of the archaeal *Haloferax* recombination occurred in tract lengths of 430Kb (20). The wide variation in recombinant tract length is likely caused by variation in DNA transfer mechanism as well as the mechanism of subsequent genomic integration.

After DNA is transferred it is processed by integration or homologous recombination machinery. These pathways further contribute to the pattern of recombination in the genome. The mechanism of genetic recombination in *S. islandicus* and other Archaeal species is not clear, with hypotheses ranging from RadA-mediated homologous recombination to a lambda-red like mechanism due to the very short recombinant tract lengths observed in transformation experiments (30,31). The results of the DNA transfer and recombination process can be analyzed by tracking linkage between selectable markers or tracking individual SNPs through genome sequencing. The size of recombinant tracts influences linkage disequilibrium and the efficiency of selection.

In natural population studies, the rate of recombination may also vary by genomic region, as we previously inferred regional rate variation from population genomic data of *S. islandicus*, with several large genomic segments exhibiting low levels of recombination (11). Variation could be due to mechanistic features of the genetic transfer and recombination processes or the result of environmental selection acting upon recombinant organisms. Here we test for the presence of recombination rate variation across the *S. islandicus* chromosome by analyzing the genome sequences of recombinants resulting from crosses of two genetically distinct strains of *S. islandicus* performed in the laboratory environment. We use multiple independent methods to calculate key parameters of recombination including recombination tract lengths, transfers of integrated plasmids, and genome-wide recombination rates.

## Results and Discussion

### Crossing divergent strains produces recombinant genotypes with clear recombination between the selected markers

To generate recombinant strains of *S. islandicus* for analysis we chose two strains isolated from a single hot spring in the Mutnovsky Volcano region of Kamchatka, Russia that vary in their core genome by about 0.2%: M.16.4 and M.16.27. We first identified a set of polymorphisms between the two parent strains (M.16.4 and M.16.27) by comparing sequencing reads from each strain against the other with breseq (Table S1). We constructed a whole-genome alignment and found 7,899 SNPs within 2.4Mb of core genome segments. We previously constructed a triple mutant of *S. islandicus* M.16.4 (RJW004: M.16.4 Δ*pyrEF* Δ*argD* Δ*lacS*). By crossing this strain with wild-type M.16.27, we selected for recombinants of both recipient backgrounds: M.16.4 Δ*pyrEF argD*^*+*^ Δ*lacS* that had incorporated the *argD* locus from M.16.27, and M.16.27 Δ*pyrEF argD*^*+*^ *lacS*^+^ that had incorporated the *pyrEF* deletion from M.16.4. The unselected marker *lacS* allowed for screening colonies’ background using X-gal staining. We isolated 133 recombinant colonies and sequenced their genomes to characterize recombination events.

We defined ‘recombinant tracts’ as any single or contiguous set of SNPs matching the donor sequence flanked on both sides by a recipient SNP. In this region of DNA between the donor DNA sequence and the recipient DNA sequence exists a recombination breakpoint. Because the mean separation between SNPs is approximately 300bp genome-wide, this method may overestimate the lengths of shorter recombinant tracts when multiple events occur over this distance, i.e. switching from donor to recipient and back to donor sequence without any polymorphisms to detect these changes. This method also likely overestimates the sizes of short recombinant tracts since recombination breakpoints could have occurred anywhere in the intervening space between the called donor and recipient SNPs, but we calculate it as the full length between the recipient SNPs.

We identified, as expected, recombination breakpoints between the selected markers *pyrEF* and *argD* in all 133 sequenced colonies (Figure 2). We called recombinant tracts that included the selected markers ‘selected tracts’. These selected tracts varied greatly between the two backgrounds, with long tracts associated with incorporation of the *pyrEF* deletion into M.16.27 and much shorter tracts involved in the insertion of *argD* into RJW004 (Fig. 2B). The shorter tracts found in insertions of *argD* into RJW004 are predominantly characterized by insertion of only the selected marker with no surrounding donor DNA (small gray boxes in Figure 2). When these tracts are excluded from the analysis, there is no significant difference between the selected tract lengths in the two backgrounds (p-value here).

**Figure 1.**
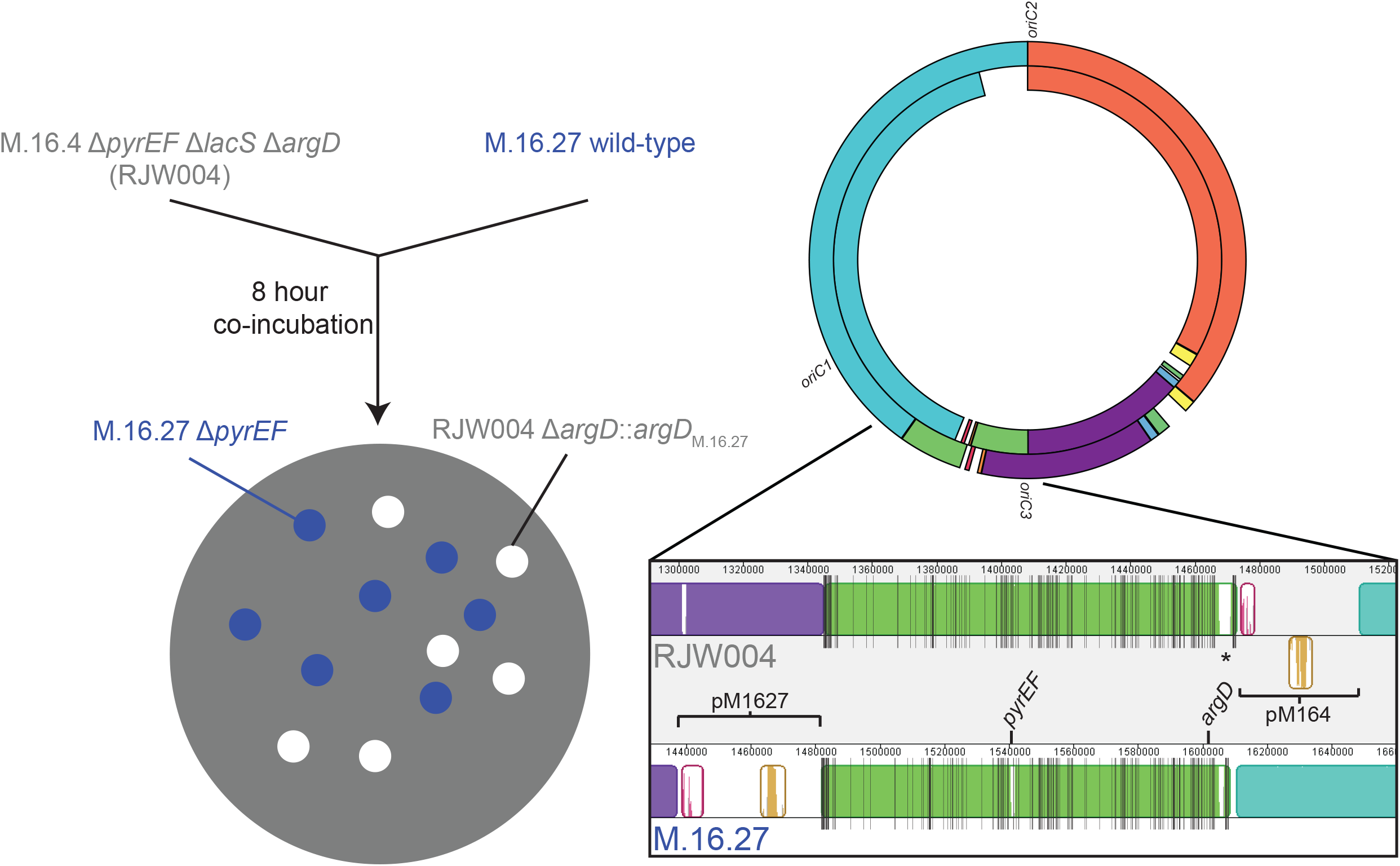
Experimental design and alignment of the parent genotypes. Strains were mixed for 8 hours, followed by plating on selective media. B) Circos plot of a Mauve alignment of M.16.27 (outer) and RJW004 (inner). Local colinear blocks (LCBs) with the same color are contiguous and homologous between strains, and blank spaces are regions with no homology between strains. The three origins of replication are marked. The zoom-in represents the alignment of the region containing the selected markers. Individual vertical lines represent the positions of SNPs between the two strains. pM164 and pM1627 are integrated conjugative plasmids.

**Figure 2.**
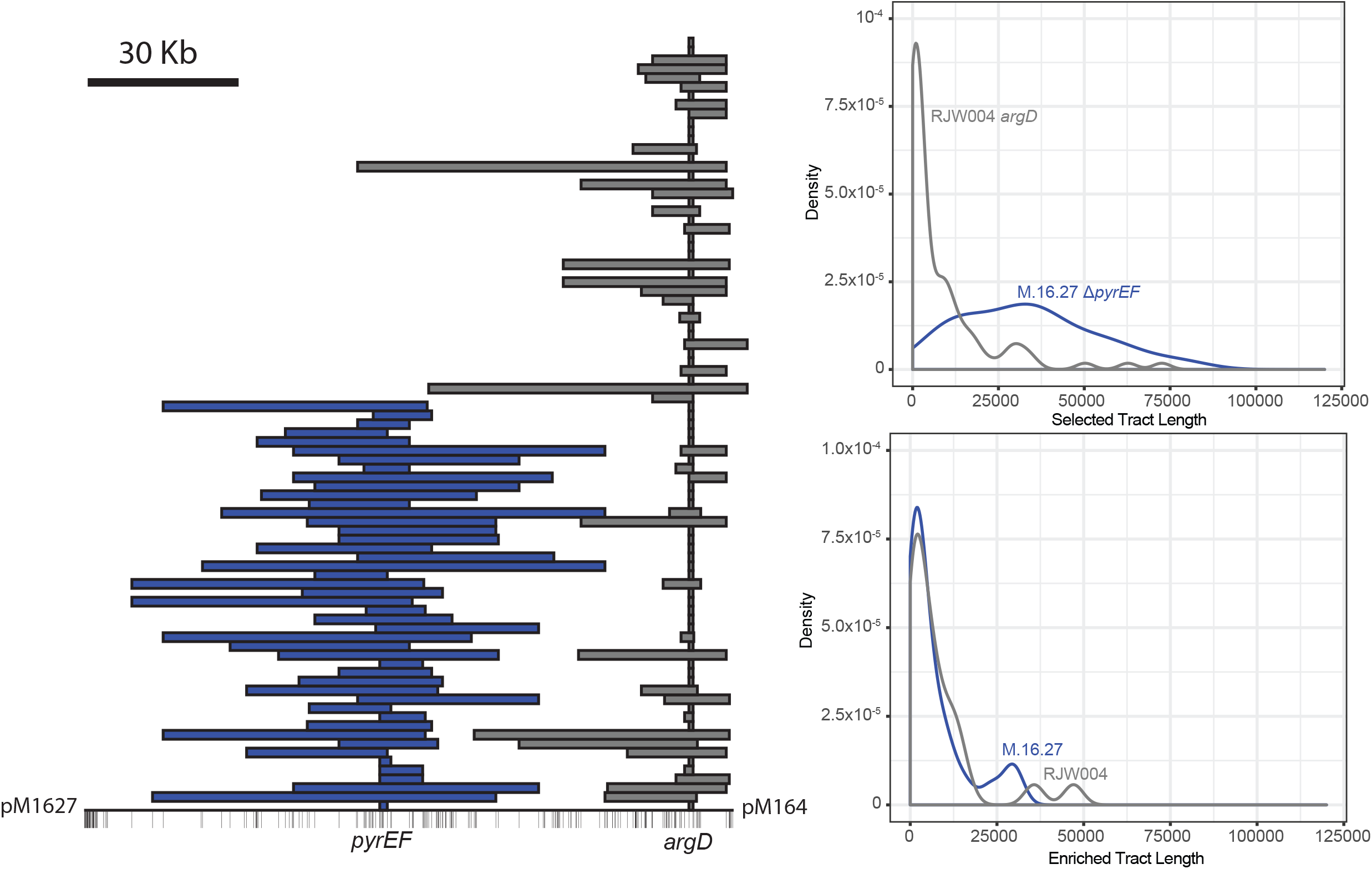
Recombination events near the selected markers. A) The conserved genomic segment containing the selected markers. Vertical lines represent SNPs (RJW004-background coordinates). Recombinant tracts in M.16.27-background strain are shown in blue, and tracts in RJW004 are shown in gray. Recombination in M16.27 contains Δ*pyrEF*, while recombination in RJW004 contains *argD*. B) Frequency distributions of selected tracts containing the marker and C) enriched tracts occurring within the selected marker region, but not containing the marker. M.16.27-background events shown in blue, RJW004-background events are shown in gray.

The *pyrEF* locus has a greater size of homologous DNA surrounding it, which may enable longer tract incorporation. We called recombinant tracts occurring on the same locally collinear block as the selected markers, but not directly including the marker ‘enriched tracts.’ These tracts likely belonged to the same fragment of donor DNA during genetic exchange but then became unlinked from the selected locus during subsequent recombination.

Both strains M.16.27 and M.16.4 have integrated plasmids in different regions of the chromosome around the *pyrEF* and *argD* markers. We identified recombination tracts in the chromosomal copies of each plasmid in both recombinant backgrounds (Figure 2C). This suggests that both plasmids can transfer between strains and may contribute to recombination frequencies in these regions.

### Recombination events often occur outside the selected markers with short tract lengths

A previous study in *S. acidocaldarius* found evidence for recombination events unlinking non-selected markers after genetic crosses (24) but could not investigate their precise position or length. While ‘enriched tracts’ qualified as non-selected recombination events, their proximity to the selected markers may have enabled their enrichment by being carried on the same piece of DNA as the selected tract. Furthermore, the selected markers in these experiments are near an integrated plasmid which may be involved in DNA mobilization either for the region surrounding the selectable markers or for the whole chromosome (27). We instead searched for truly non-selected recombination events throughout the chromosome, and we identified a total of 32 such tracts in 10 M.16.27-background colonies and 115 tracts in 31 RJW004-background colonies (Figure 3).

**Figure 3.**
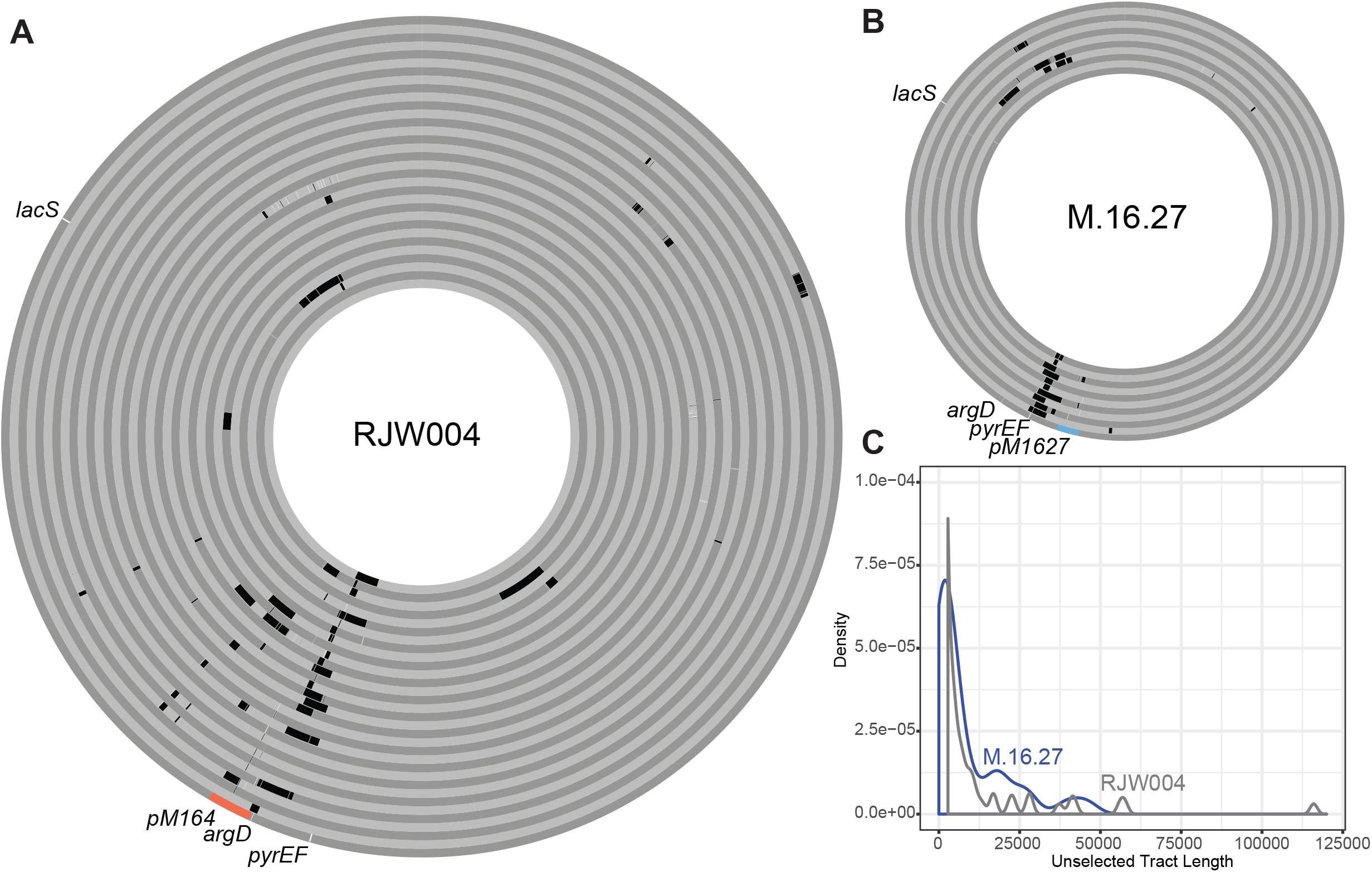
Distribution of unselected recombination events. A) Tracts were plotted in Circos, and each concentric circle corresponds to an individual recombinant colony in the RJW004 background. B) Same as (A) in the M.16.27 background. C) Frequency distribution of unselected tracts.

The length distributions between the two backgrounds were not significantly different (Wilcoxon rank sum test, *p* = 0.092), so we combined the two distributions for further analyses. The combined distribution was significantly right-skewed (Shapiro-Wilk test, W = 0.49, *p* < 2.2x10^-16^), with a mean of 6.7Kb and a median of 1.5Kb. The short length of these tracts raised issues related to their detection. We could only identify tracts that incorporated one or more donor SNPs, and their lengths were conservatively calculated as the distance between adjacent reference-matching SNPs. Using the distribution of SNPs across the genome, we simulated recombinant tracts for a subset of the observed tract lengths to estimate the frequency of missing tracts as well as the accuracy of our length measurements. The simulation suggested that we lacked the power to detect very short tracts, with the shortest tract experimentally measured (40bp) being missed 90% of the time, and only tracts longer than about 3,000bp being detected with greater than 90% efficiency (Figure S1). Applying a correction based on this result significantly shifted the frequency distribution toward shorter tracts (Wilcoxon rank sum test, *p* = 3.4x10^-7^, Figure S2), with a median of 432bp. Even when we positively identified the short tracts in the simulation, their measured lengths were overcalculated by our conservative methods at the shorter end of the distribution (Figure S3). When we considered short tracts as all unselected measured tract lengths shorter than 10,000bp, we found that this distribution was not significantly different from the one generated from simulated tracts of 360bp (Wilcoxon rank sum test, *p* = 0.55). These two results suggest that the detection bias against short tracts and overestimation of measured lengths have obscured the true distribution of tracts, and the true median may be even less than our 432bp estimate.

Short and often discontinuous recombinant tracts had been experimentally observed in the related archaeon *S. acidocaldarius* in regions within and surrounding the selectable marker *pyrE* (33), and they had been previously inferred from population genomics studies of *S. islandicus* (11,32). However, these results are the first direct observations of short-tract recombination occurring outside a selectable marker in experimental data. Short patch recombination has been observed in *S. cerevisiae*, with a higher frequency found in *msh2* mutants that lack long-patch excision repair (34). Short patch recombination is also evident in transformations of bacteria such as *S. pneumonia, H. influenzae*, and *H. pylori* (17,19,35). While *Sulfolobus* species do not contain homologs of MutS or MutL mismatch repair proteins, they do have a the NucS/EndoMS protein that could be involved in mismatch repair, although it is not yet known whether this pathway can lead to short tract repair (36). In contrast, the less frequently observed long tracts have properties similar to crossover events in eukaryotes, as well as the results of conjugations-based mechanisms in *S. agalactiae* and *M. smegmatis* (18,29). Recombination initiated at double-strand breaks as the result of homing endonuclease activity in polyploid halophilic archaea have observed recombination events as long as 50Kb (37), indicating that large pieces of DNA can move during genetic transfer in archaea. Because of the enrichment of discontinuous recombinant tracts near the selected markers, as well as the clustering of small patches of discontinuous tracts elsewhere in the recombinant genomes, we believe that large pieces of DNA, perhaps even the whole chromosome are transferred, but the recombination mechanism leaves discontinuous patches of donor DNA. It is not clear from these data whether this occurs through conjugation, cell fusion, or represents multiple mechanisms for homologous gene transfer in *Sulfolobus*.

### MARBL, a method for identifying recombination breakpoints in deeply sequenced large colony pools

Sequencing individual recombinant genotypes offered insight into parameters of recombination at the individual level, namely the frequency and sizes of recombinant tracts that arose from recombination. However, we also sought to understand whether recombination rates varied around the chromosome, which required observing enough events to cover the chromosome with an expectation of observing multiple events at any position. To attempt this, we sequenced several pools of recombinant colonies, with each pool containing many colonies of a single genetic background (Table 1). Although we could not separate individual recombination events with this method, we calculated donor SNP frequency throughout the genome. The pools ranged in size from 79 to 9,026 colonies, with per-colony sequence depths ranging from 1,880x to 18,300x.

We confidently measured variation in donor SNP frequencies directly surrounding the selected markers, showing a pattern of decreasing donor SNP frequency with increasing distance from the selected marker characteristic of recombination experiments including those in *S. acidocaldarius* (33). Elsewhere in the genome, donor SNP frequencies could not be calculated confidently above the rate of sequencing error, in part due to the inability to obtain totally pure colony pools by X-gal staining.

We next developed a method for calculating position-specific recombination rates directly from sequence data without needing to keep pools of a single background genotype. We named this tool MARBL, an acronym for <u>M</u>ultiply-<u>A</u>nchored <u>R</u>ecombination <u>B</u>reakpoint <u>L</u>ocator (see methods). MARBL works by using multiple SNP calls on either side of a putative recombination breakpoint, thereby reducing the influence of sequencing error (see Methods; equation 1). MARBL identifies breakpoints in paired-end reads that contain four or more polymorphic sites and calculates a per-base-pair recombination rate for the intervening positions (Figure 5).

**Figure 4.**
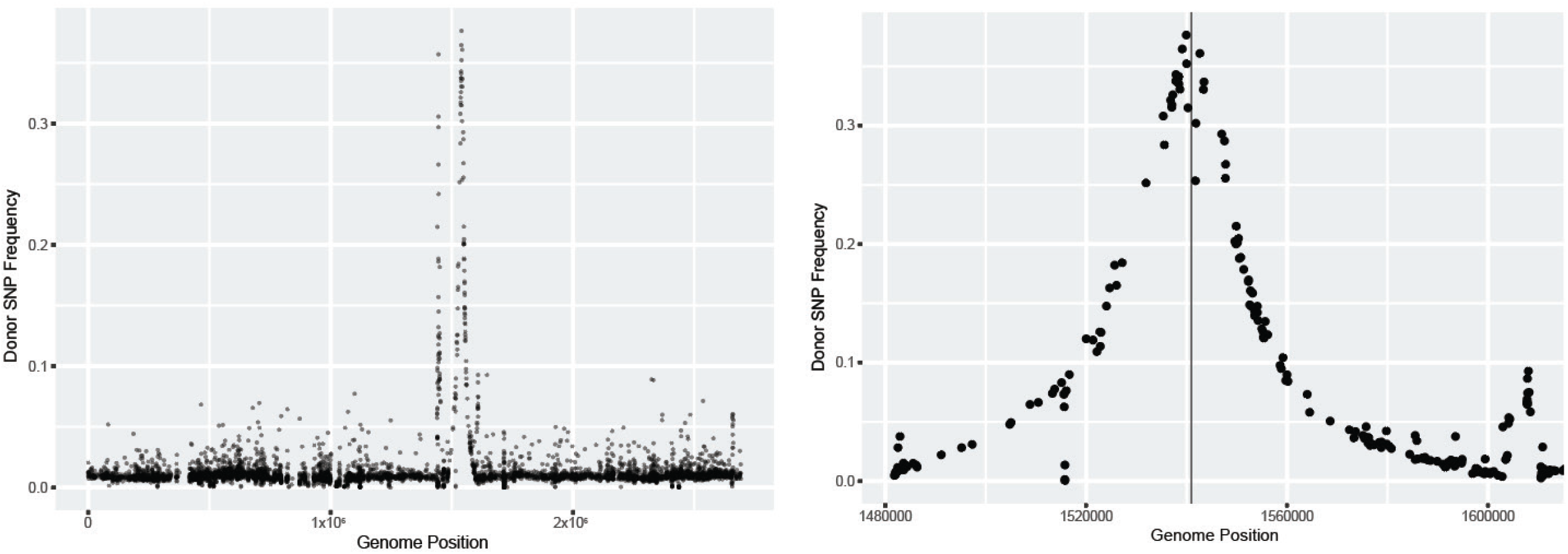
Donor SNP frequencies calculated from a deeply sequenced single-background colony pool. A) Donor SNP frequencies were calculated at each polymorphic site, and values were potted on M.16.27-background coordinates. B) Zoom-in view of the region surrounding the selected marker. Δ*pyrEF* is represented as a vertical bar. The left and right bounds are the ends of the conserved genomic segment containing the selected markers.

**Figure 5.**
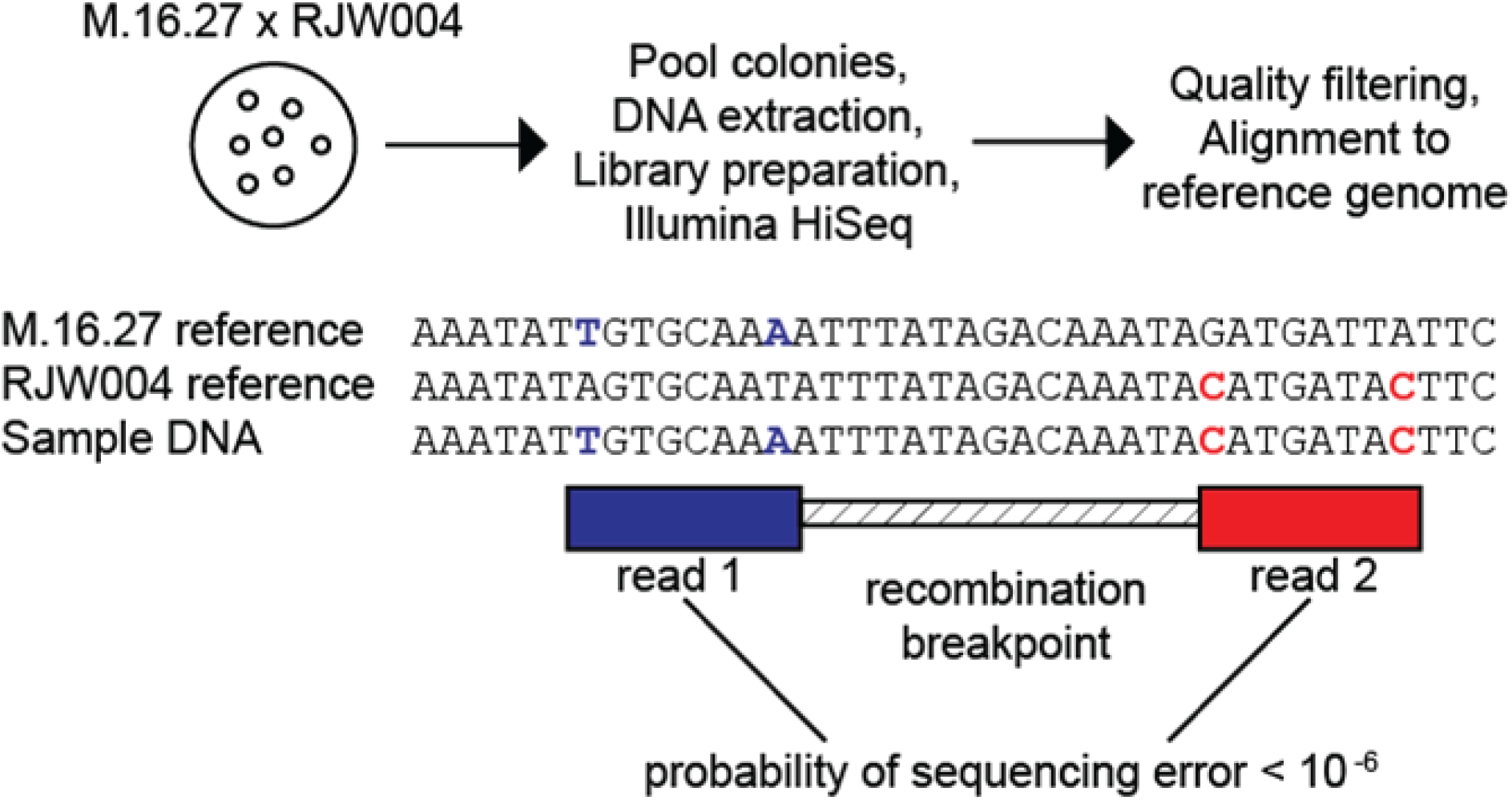
Summary of the MARBL workflow. DNA is extracted from recombinant colony pools and sequenced, followed by quality control and alignment to a reference. MARBL then finds crossovers between the two genotypes that pass a statistical significance threshold.

We ran MARBL on sequencing reads obtained from deep sequencing of genomic DNA from the two individual parent strains M.16.27 and RJW004, and we identified no recombination, indicating that MARBL is not prone to false detection due to sequencing errors. We also tested MARBL against mock recombinant reads generated using bbmap (see Methods). We found that MARBL did sometimes lack power to detect true recombination events, likely due to complexities of the M.16.27-RJW004 alignment in certain genomic regions, especially those containing repetitive sites. We removed these potential junctions where MARBL failed to detect recombination from further analysis. We further ran MARBL on a mixed gDNA sample of M.16.27 and RJW004, to identify whether MARBL detected any artefactual recombination from library preparation or the sequencing process. We did identify 218 false breakpoints over a total of 3.4 million total SNP pairs with covered read pairs, giving a false positive rate of 6.4x10^-5^ events per test.

To obtain a dataset rich in recombination breakpoints, we performed two additional large-scale genetic crosses of M.16.27 x RJW004, collecting 18,767 colonies and 20,131 colonies respectively. We sequenced these colony pools to depths of 22,800x and 66,000x respectively per base-pair, corresponding to a mean depth of 1.2x and 3.3x per base-pair for each colony. We also ran MARBL on the large, deeply sequenced B1 colony pool (Table 1). MARBL identified recombination breakpoints in all three colony pools at rates 3-10 times higher than the false positive rate (Figure 6). We also found no significant correlation between recombination rates calculated from the chimeric DNA control run and the recombinant colony pool runs, indicating that while false positives may be inherent to the data, we found no evidence for false positives driving the patterns that we found (R=0.037, *p* = 0.36).

**Figure 6.**
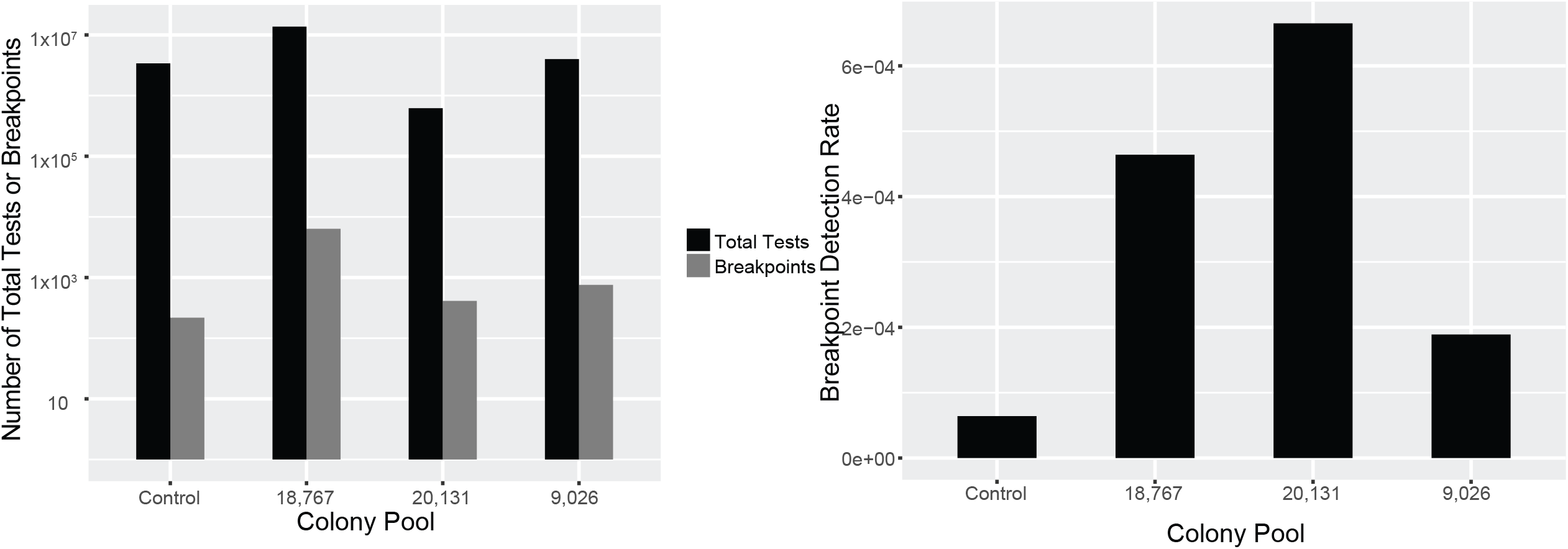
Statistics of MARBL run on several colony pools, including a false positive control. A) The absolute number of tests for recombination breakpoints, as well as the total number of breakpoints are shown. B) The detection rates shown were calculated as the number of breakpoints divided by the total number of tests from (A).

We calculated recombination rates for each position in the genome for each of the colony pools using MARBL (Figure 7). The weighted recombination rates showed peak values around the selected markers as well as in the integrated plasmids, in line with previous observations of recombination in the individual colony sequences as well as the partitioned colony pools (Figure 7). Outside of the plasmid and selected markers, recombination rates were not found to show statistically significant variation around the chromosome according to Mantel tests for spatial auto-correlation between recombination rate differences and positional distances among measurement sites, k-means clustering to determine if outlier data-points are clustered in particular genomic locations, and ANOVA of binned recombination rates at various bin sizes (see Methods, Table S3).

**Figure 7.**
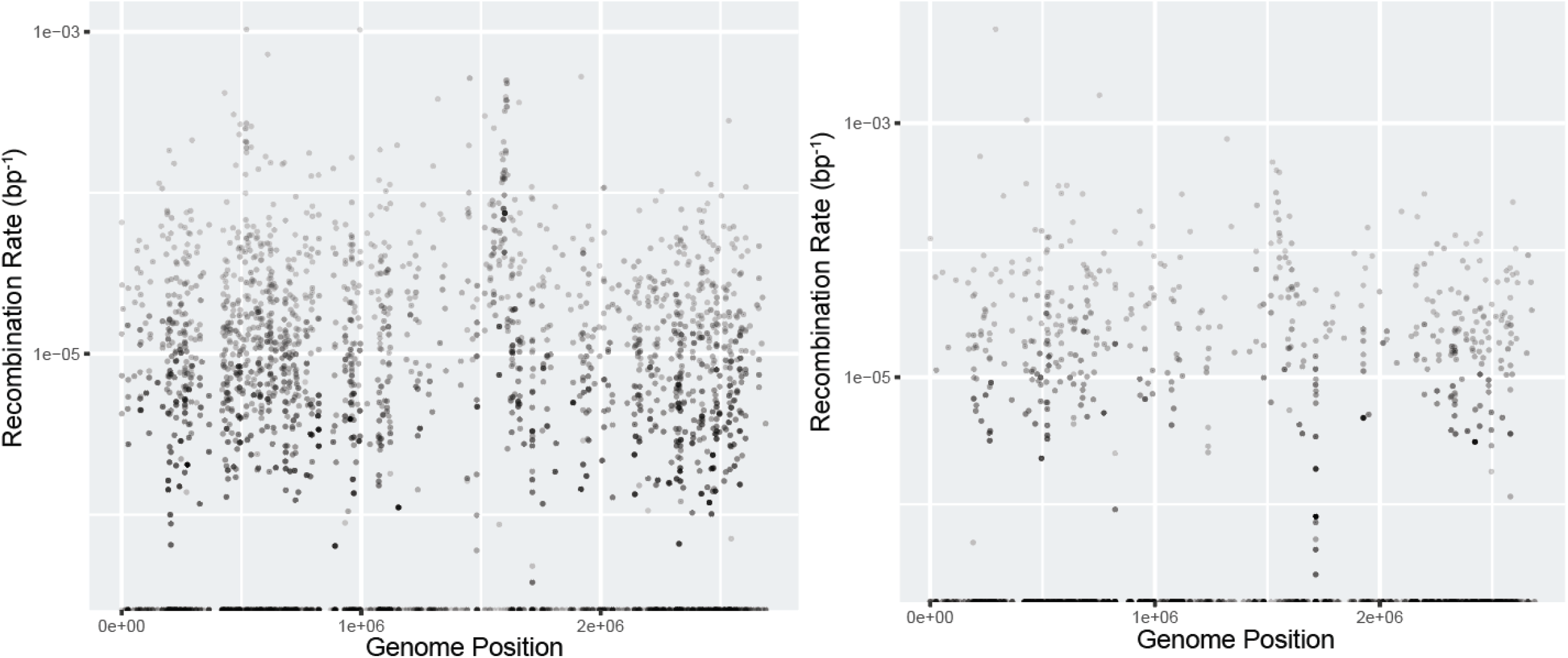
Genome-wide recombination rates estimated by MARBL for deeply sequenced large colony pools. A) The 18,767-colony pool. B) The 9,026-colony pool. The position of each point was calculated as the midpoint between the two adjacent SNPs over which recombination rates were calculated. Points were shaded by their weight, as calculated by the product of the number of tests and the length of the sites, which was chosen as a metric corresponding to the expected number of detectable recombination breakpoints.

MARBL analysis of the large colony pools, as well as the six smaller sorted pools identified remarkable consistency in mean genome-wide recombination rates. The sorted M.16.27-background colony pools contained significant levels of spontaneous *pyrEF* mutations, calculated as the presence of DNA covering regions of the genes that should have been deleted in true recombinant colonies. Since these spontaneous mutants likely did not recombine during the experiment, the recombination rates in these pools were corrected according to the spontaneous mutant ratio (Table S2). The seven recombinant colony pools had a mean recombination rate of 2.9x10^-6^ bp^-1^ ± 8.4 x 10^-7^ bp^-1^ (S.D.). All the colony pools fell within the 99% confidence interval of this mean, while the chimeric mixture control pool had a rate that fell 3.2 standard deviations below this mean, indicating that the colony pools generally showed similar levels of genome-wide recombination, significantly above the false positive background rate.

The finding of constant genome-wide recombination rates is in contrast with previous findings of low-recombination zones (LRZs) in natural population data of *S. islandicus* (11). Recombination rates within the LRZs in the 18,767-colony pool were not significantly different from the calculated genome-wide recombination rate: approximately 2.7 x 10^-6^ ± 2.1 x 10^-6^ bp-1 (95% C.I., unadjusted weighted mean). Such a contrast between population studies and laboratory measurements implies that selection acting upon recombinant genotypes in nature has played a role in defining those previously described patterns. One possible explanation for identifying recombination rate variation in nature, but not in the laboratory, is selection for or against specific recombinant genotypes in nature.

By combining the power of colony pooling and deep sequencing to identify genome-wide patterns and high-resolution views of highly recombining genomic regions with precise discrete recombinant tract estimation of individual colony sequencing, we have presented an overview of parameter estimates for recombination in *S. islandicus*. While previous studies have assessed various parameters at a single locus, this work identifies recombination occurring throughout the genome, gaining information about how much of the genome is accessible to the recombination machinery during DNA exchange. We found that short tract recombination is likely highly prevalent throughout the *S. islandicus* genome, which in principle can remove linkage between closely spaced sites over the course of a single recombination event. This has implications for *S. islandicus* population biology, as selective sweeps may not leave strong patterns of linkage disequilibrium if recombination events can break linkage over hundreds of base pairs. Finally, we did not identify variation in recombination rates among genomic regions, suggesting that patterns observed in nature are shaped by environmental selection, rather than inherent recombination rate variation throughout the *S. islandicus* chromosome. However, these results are also dependent upon laboratory observations of cell-cell mating experiments. If other mechanisms of gene transfer are more common in nature, such as plasmid conjugation or viral transduction, recombination rates may vary from what we have observed with cell-cell matings.

## Methods

### Genetic cross assay

Cultures of M.16.27 (wild-type) and RJW004 (M.16.04 Δ*pyrEF*Δ*argD*Δ*lacS*) were grown to OD 0.3 in dextrin-tryptone liquid medium (32,38) supplemented with 20µg/ml uracil (Sigma-Aldrich) and 20µg/ml agmatine (Sigma-Aldrich), then mixed at equal ratios for co-incubation at 78°C for 8 hours, with shaking at 180 rpm. The cell mixture was then washed three times in medium lacking agmatine to remove residual agmatine, followed by plating and beta-galactosidase staining on both selective dextrin-tryptone plates containing 50µg/ml 5’-FOA (Toronto Research Chemicals) and 20µg/ml uracil and non-selective plates containing 20μg/ml uracil and 20μg/ml agmatine.

### Colony pooling and DNA extraction

After 10 days incubation of plates at 78°C, colonies were picked from the surface of the selective plates and re-suspended in DT liquid medium supplemented with uracil at room temperature. Table 1 shows the colony pools that were obtained and analyzed. The B1-3 and W1-3 colony pools come from three independent experiments (1-3) and were stained with X-gal dissolved in DMF prior to creating separate pools based on blue/white coloration (B/W). Colony pools were pelleted and re-suspended in TE buffer pH 8.0. Liquid cultures were grown to 0.2 OD600, pelleted, and re-suspended in TE buffer pH 8.0 join here. Cells were lysed using GES [60% w/v guanidine thiocyanate (IBI Scientific), 4.2% w/v EDTA-tetrasodium (Sigma-Aldrich), 0.5% w/v N-lauroylsarcosine (Sigma-Aldrich)], neutralized by addition of 150mM ammonium acetate (Fisher Scientific), and then treated with 24:1 chloroform/isoamyl alcohol (Fisher Scientific). Phases were separated by centrifugation at 13,200 rpm for 5 minutes, followed by extraction of the aqueous phase. DNA was precipitated with isopropyl alcohol (Fisher Scientific) and washed with 70% ethanol (Decon Laboratories, Inc.) prior to resuspension in EB buffer (Tris pH 8.0).

### DNA library preparation and sequencing

A description of each DNA sample can be found in (Table 1). For the 18KCOL colony pool and CHI_MIX sample DNA was sheared using Covaris to isolate DNA fragments in the size range of 500-700bp (this size range was expected to produce reads that maximized coverage of genomic variation). This fraction was prepared for sequencing using the Hyper Kapa Library Preparation Kit, with three rounds of post-preparation PCR amplification. The resulting DNA libraries were sequenced on an Illumina HiSeq2500 for 251 cycles from each end of the fragments using a Rapid TruSeq SBS kit version 2. Fastq files were generated and demultiplexed with the bcl2fastq v.1.8.4 Conversion Software (Illumina).

For the B1-3, W1-3, M.16.27, and RJW004 samples, DNA was prepared using the Nextera XT DNA Sample Preparation Kit (Illumina). Prepared DNA was sequenced on an Illumina HiSeq2500 for 151 cycles from each end of the fragments using a Rapid TruSeq SBS kit version 1. The B1 pool was re-sequenced on another lane using the same procedures, resulting in its higher depth of coverage.

### Generation of simulated sequencing reads

Simulated sequence reads were generated using the bbmap toolkit’s randomreads.sh script . Parameters were used to mimic the libraries used in the 250bp paired-end Illumina HiSeq runs, with a fragment size distribution of 330-900bp, and total read count of 250,000,000. The M.16.27 genome can be found at NCBI Accession #NC_012632.1, and the M.16.04 genome can be found at NCBI Accession #NC_012726.1. The RJW004 genome has been sequenced previously and polymorphisms in the genome were inferred via comparison with the M.16.04 ancestor reference using breseq.

### Read trimming and quality filtering

Reads were trimmed of adapter sequence using cutadapt (39). For the 18KCOL and CHI_MIX samples, the first mate pair file used ‘AGATCGGAAGAGCAC’ and the second mate pair file used ‘AGATCGGAAGAGCGT’ as the adapter sequences. For the Nextera-prepared pools (B1-3, W1-3, M.16.27, RJW004), the adapter sequences used were ‘CTGTCTCTTATA’ for both files. Reads were quality trimmed using sickle, with a 10bp average quality score cutoff of 25 and minimum length cutoff of 50bp.

### Alignment of reads with the reference genomes

Reads were aligned to a reference genome using Novoalign v.3.02.12 and a license provided by the University of Illinois HPC-Bio group. To avoid soft-clipping in alignments, the option for a full Needleman-wunsch alignment was invoked. Output file format was required to be a .sam file for compliance and functionality with downstream tools.

### SNP database creation

Mock reads from M.16.27 and RJW004 were aligned to the M.16.27 reference genome using novoalign. Alignments in the SAM file were analyzed for variants at each position. Positions with 100% consensus in both genomes were used for downstream analysis, while positions containing within-sample conflicting calls were discarded. The M.16.27 and RJW004 genomes were also aligned using Mauve v.2.3.1 with default parameters. Only SNPs detected using Mauve and mock read alignment were kept for the SNP database. Of the kept positions, those with M.16.27 and RJW004 calling the same base were used to calibrate error rates as well as estimate mutation rates, while those that had different calls between genomes were used as the SNP database to identify recombination breakpoints.

### Sample-specific error rate calibration

The statistical significance of breakpoint identification is determined by a likelihood of sequencing error deduced from internally-derived calibration of error rate from each sample using the database of positions identical between M.16.27 and RJW004. Briefly, every base call is binned according to multiple parameters, including quality score, position within the read, and preceding called dinucleotide. The error rates corresponding to each parameter value are then calculated using the frequency of reference matches and mismatches in the sample itself using in-house perl scripts. Next, for each combination of nucleotide call, preceding dinucleotide, and quality score, variation in error rate over all positions within the read was interpolated using polynomial regression in R. Interpolated values were capped at a minimum of 1.1x10^-6^ and a maximum of 0.1, to avoid downstream software errors, as well as ensure that no single nucleotide call passes the significance threshold for recombination detection. The first 5 5’-end nucleotides and the last 2 3’-end nucleotides were removed from analysis due to irregular error rates for 250-bp reads, while the first 2 5’-end nucleotides were removed for the 150-bp reads.

### MARBL, Multiply-Anchored Recombination Breakpoint Locator

The sample alignment file is then searched for paired-reads that cover multiple SNPs according to the SNP database file. Those paired-reads containing multiple SNPs are then tested for the potential to yield statistically significant recombination breakpoints (p < 1x10^-6^). This significance threshold was decided on the basis of maximizing the detection rate while limiting the influence of false positives (Figure S4). If the sum of the products of error likelihoods on either side of a candidate junction is less than 10^-6^, then this junction contains information that could lead to detection of a recombination breakpoint and is counted as a test. For any junction in any given read-pair, the test metric is calculated as the maximum theoretical number of breakpoints that the read-pair can identify divided by the number of test-able junctions in the read-pair. The strain origin of the DNA (M.16.27 or RJW004) is recorded for each SNP, and any junction that contains ≥2 SNPs of one genotype on one side, and ≥2 SNPs of the other genotype on the opposite side is determined to be the site of a recombination breakpoint with a value of 1.

### Detection of recombination breakpoints in simulated recombinant sequences

First, 5Kb sequence chunks were extracted from the M.16.27 reference genome, centered on each of the detected sites in the 18K_colony sample MARBL output. Next, all polymorphic sites on either the left or right side (both were done) of the recombinant junction were substituted with the RJW004 polymorphism. Then, bbmap’s randomreads.sh was used to generate 10,000 paired-end 250bp reads for the recombinant 5Kb sequence chunk. These reads were subjected to the same treatment as above (with the exception of quality filtering as there are no errors), and run through MARBL. Junctions that showed zero recombination from one or both sides were removed from further analysis to avoid false negative results.

### Statistical analysis of recombination rate variation

All statistical analyses were performed on the 18,767-colony pool. The Mantel test was used to test for correlation between recombination rate differences and positional distance among measured sites. With the selected marker included, there is a significant correlation between the genome proximity and recombination rate (r=0.03, *p*=0.001). Without the selected marker region, but including the integrated plasmid, the magnitude of the correlation decreases, but is still significant (r=0.011, *p*=0.011). Without the integrated plasmid measurements, this correlation becomes non-significant (r=0.0075, *p*=0.059), despite the fact that a subtle auto-correlation exists for distances <500bp due to the possibility of analyzing the same recombination breakpoint multiple times at slightly different measurement positions.

K-means clustering was used to compare the clustering of positions of outlier measurements to the clustering of random measurements. Within-cluster variance was calculated for 1,000 similarly-weighted replicates, and the *p*-value is represented as the proportion of random measurements with less within-cluster variance than the sample. With the selected marker included, the top ten to twenty outlier loci are heavily clustered in this region (*p*<0.001). However, after the removal of the selected marker, no significant clustering is observed (Table S3).

For ANOVA, various binning sizes were used to break the genome into equal parts. unlike the Mantel and k-means testing, which test for variation over continuous scales, binning imparts a subjective assessment of “regional” variation. Therefore, multiple bin sizes were used to break the genome into variably scaled regions. While ANOVA shows significant variation between regions at the 25Kb (F=4.3; df=94, 5211; *p*<2x10^-16^) and 10Kb sizes (F=2.9; df=182, 5,123; *p*<2x10^-16^), a Tukey’s HSD test determined that all significant differences (*p*<0.05) are caused by a single bin within the integrated plasmid. Removal of the integrated plasmid from analysis shows no significant individual bin comparisons at any tested bin size from 10Kb to 250Kb.

### Running MARBL on sample data

MARBL can be accessed, along with sample datasets and usage instructions via the github repository system using the following website: https://github.com/krausedk/marbl_scripts. Data are available from the authors upon request.

## Acknowledgments

The authors thank Dr. Alvaro Hernandez and Dr. Chris Wright for their contributions in designing the DNA library preparation protocol and performing the DNA sequencing. The authors would also like to thank Prof. Sergei Maslov and Prof. Alexander Lipka for helpful comments during project design and data analysis. This work was supported by a grant to Dr. Rachel J. Whitaker from the National Science Foundation: DEB #1355171 and from NASA: NNX09AM92G.

