## Supplemental Figures for "Analyses of Genome-Wide Recombination in *Sulfolobus islandicus*"

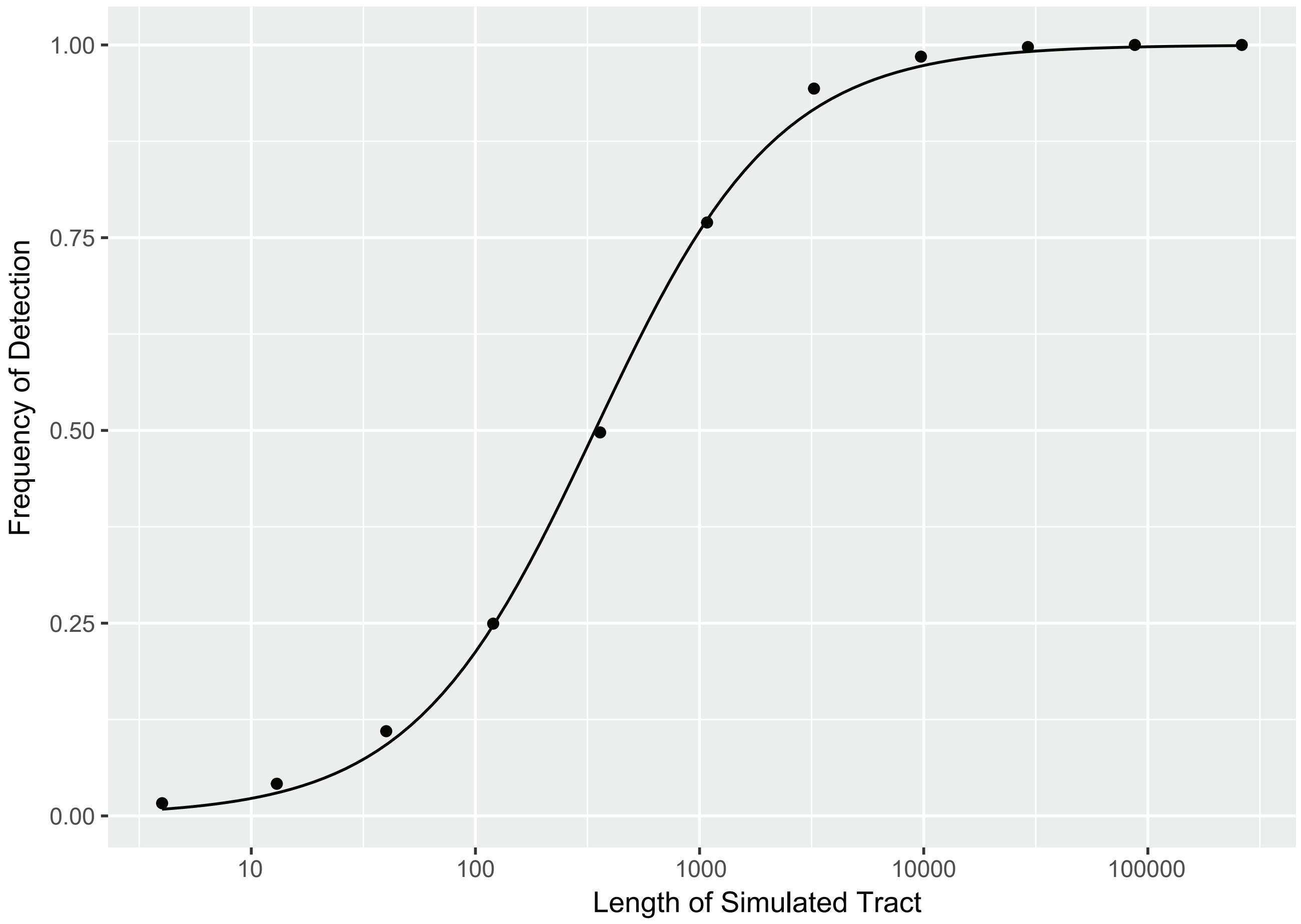


Figure S1 – Likelihood of detecting recombinant tracts is dependent upon length. Results shown are from simulations of 10,000 tracts for each length, with detection frequency measured as how often the tracts overlapped with a SNP.

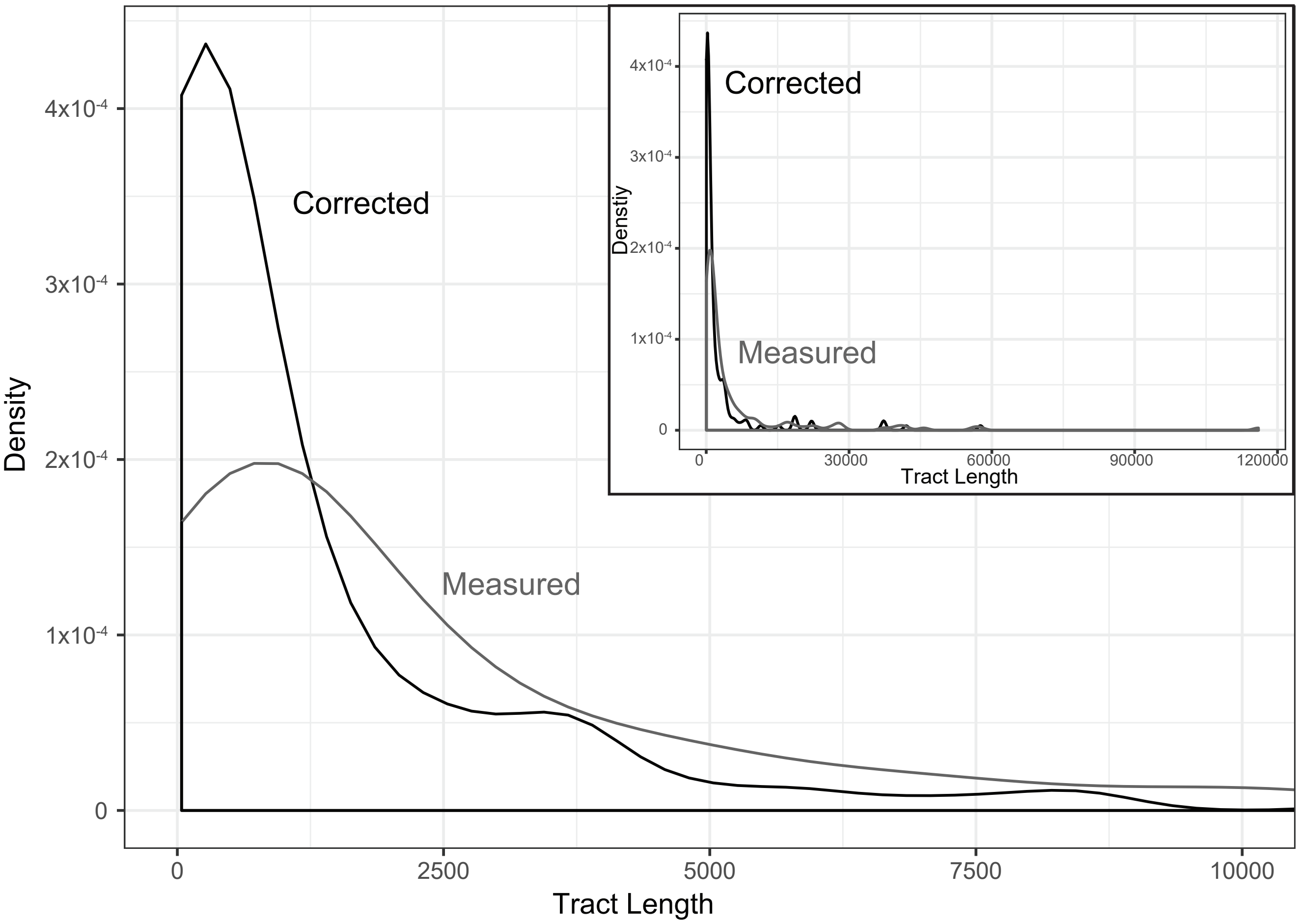


Figure S2 – Correcting the tract length distribution. The detection rates from Figure S1 were used to correct the frequency of the measured tract lengths within the initial distribution.

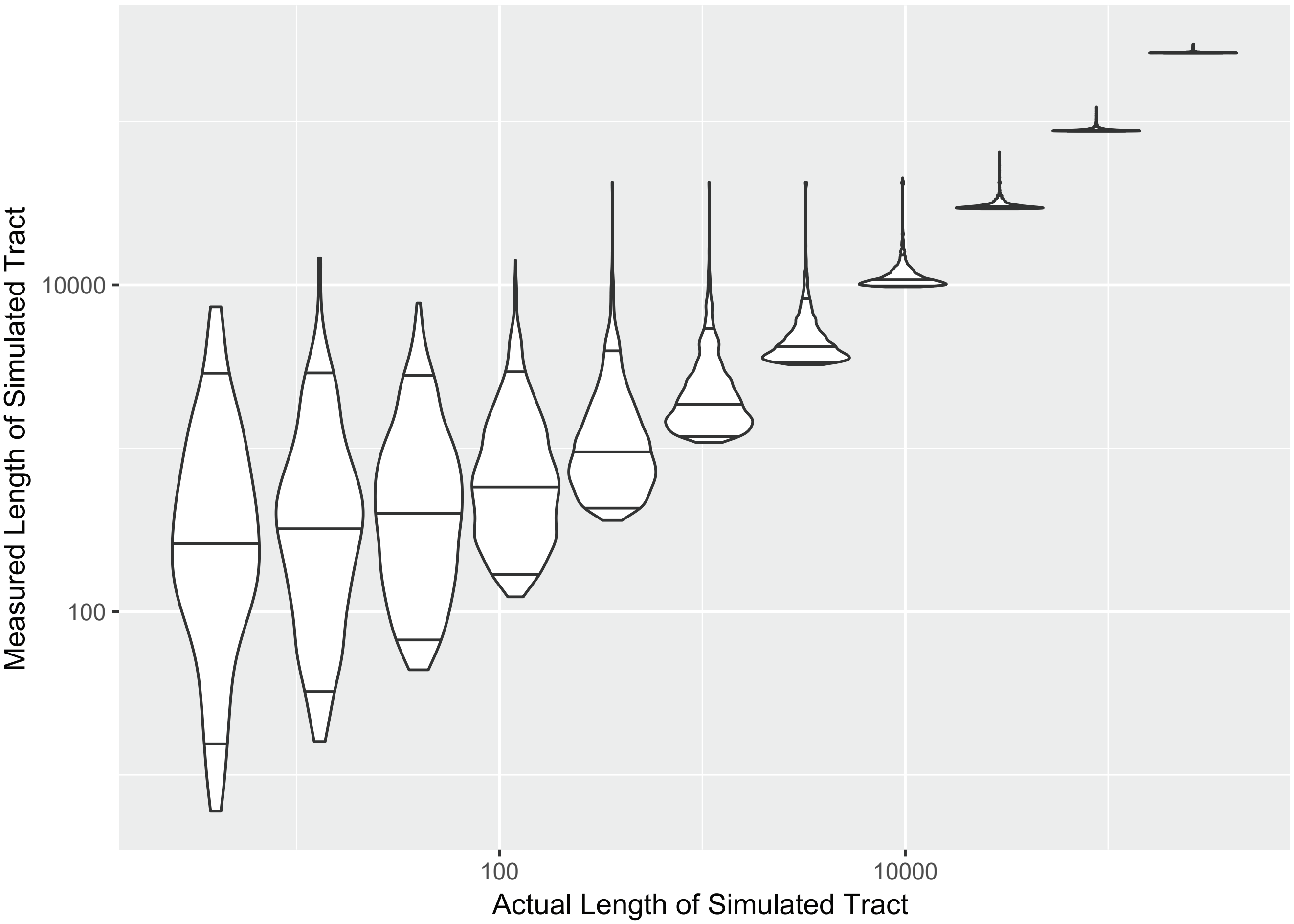


Figure S3 – Overmeasurement of tract lengths. For tracts detected in the 10,000-tract simulation, the tract length was measured according to the Methods. The tract length distribution for each of the varying simulated lengths is shown in the violin plots. Horizontal lines within the violin plots correspond to the 5%, 50%, and 95% quantiles.

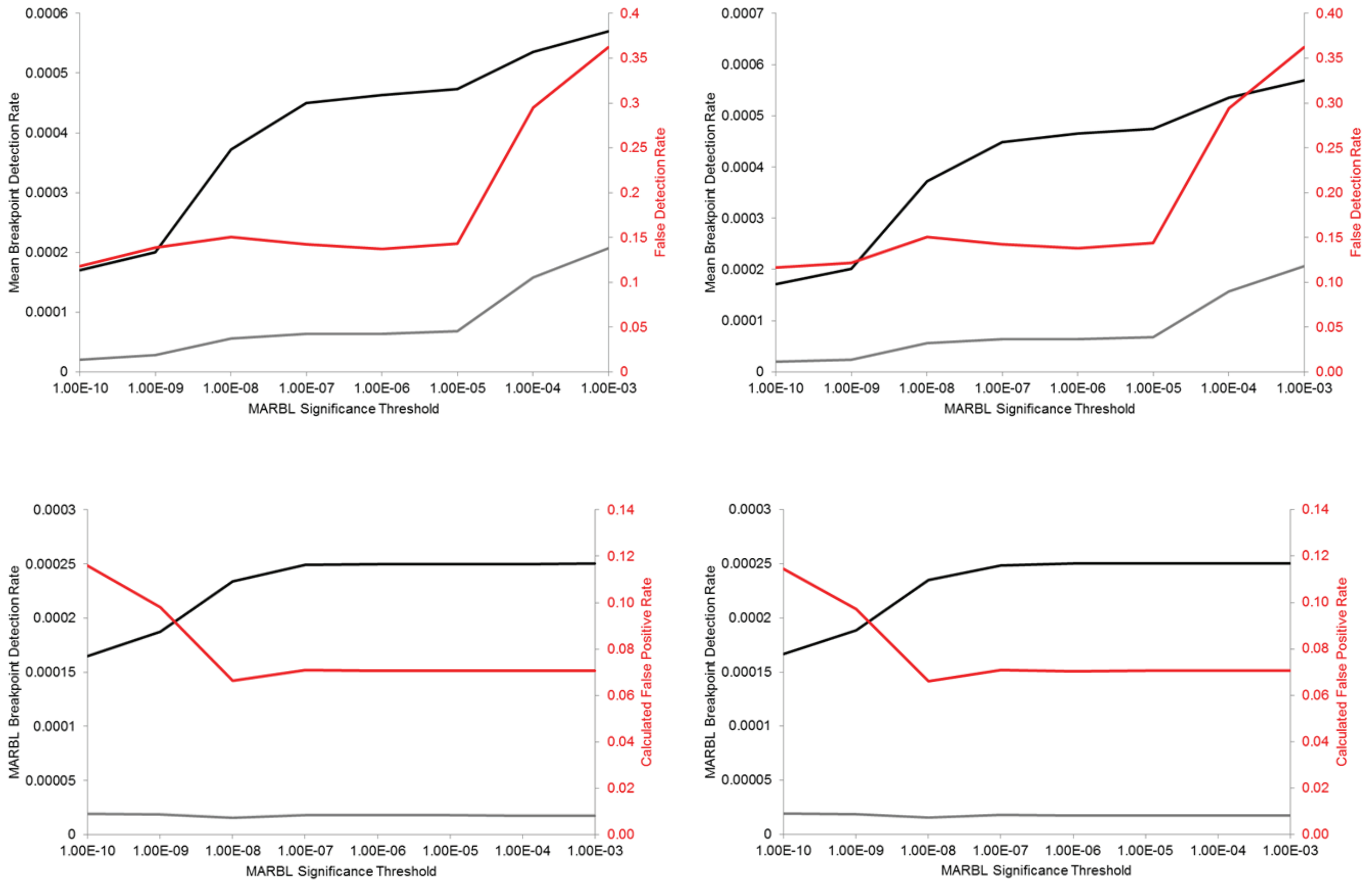


Figure S4 - Altering the significance threshold of MARBL shows that false detection is minimalized at 10-15% for p-value thresholds less than 1x10$^{-4}$. Values between 1x10$^{-7}$ - 1x10$^{-5}$ show very little variation, given a per-base-call cutoff of A) 0.001 or B) 0.01. Including a requirement for 2 SNPs to be called on either side of a junction creates the patterns found in C) and D) at 0.001 and 0.01 per-base-call-cutoffs, respectively, dropping the false positive rate to 7%.
